# SpaceBender: Denoising Spatial Transcriptomics Data to Enhance Biological Signals

**DOI:** 10.64898/2026.04.20.719715

**Authors:** Daniel G. Chen, Antoni Ribas, Katie M. Campbell

## Abstract

Spatial transcriptomics (ST) allows for the simultaneous profiling of cell phenotype (e.g. transcriptome) and physical position. Although ST data has brought about numerous new biological insights, it remains limited by noise, largely in the form of RNA diffusion. Here, we introduce SpaceBender which leverages spatial-specific information (e.g. spatial ambient RNA niches) to build upon single-cell denoising strategies. SpaceBender outperforms current ST denoising methods in simulations and *in vivo* chimeric tissues. Through case studies, we demonstrate how SpaceBender unveils hidden biological insights and increases the significance of said insights as evaluated by statistical testing. Finally, we reveal how SpaceBender may also be applied to subcellular resolution data where it removes off-target expression of neighboring cell type specific marker genes. In all, we present SpaceBender as an ST denoising method, freely available as an open-source package, that may enhance the insights the field may draw from various ST data types.

## Introduction

Spatial transcriptomics (ST) has revolutionized our ability to interrogate human biology through its simultaneous measurement of both phenotype (e.g. transcriptome) and physical position^1–3^. ST technologies have varied in their implementation; they span spot-based (i.e. multi-cellular) and sub-cellular resolutions and measurements via sequencing (e.g. SlideSeq and Visium) and imaging (e.g. Xenium, MERFISH, and CosMx). Although ST methods have markedly moved the field forward^4,5^, certain technical limitations restrain our ability to fully take advantage of this technology. For example, ST data have region-level artifacts (e.g. degraded tissue fragments) that cannot be addressed by previous single-cell methods and thus warrant the development of new computational frameworks, such as SpotSweeper^6^.

A principal limitation of ST data is RNA diffusion, whereby RNA molecules disperse from their original spatial location through random (Brownian) motion^7,8^. This leads to the incorrect quantification of gene expression in spots or cells due to the migration of RNA molecules from nearby regions. Several technologies and computational frameworks have sought to rectify this limitation by enhancing the accuracy of RNA capture^9^ and computational correction of said diffusion^7,10^. Although technological innovations will eventually overcome this barrier, much data has already been generated with ST technologies that possess, to varying degrees, RNA diffusion^11^. Thus, there is significant interest in the development of computational methods that reduce technical noise in ST data and enhance the detection of true biological signal.

Notably, methods to denoise ambient (diffused) RNA for single-cell technologies have already been well established, e.g. SoupX^12^ and CellBender^13^. Recent works have highlighted the value of adapting extra-disciplinary methods for computational biology, e.g. applying ecological methods to spatial biology problems^14^. Similarly, we propose adapting deep generative models created for single-cell denoising to ST data through several key changes that leverage spatial-specific information. Here, we demonstrate how, 1) this adaptation, which we term SpaceBender, serves as a leading computational framework to denoise ST data, 2) reveal how SpaceBender enhances our ability to draw biological insights from ST data, through case studies across physiological contexts, and 3) how SpaceBender can not only be applied to spot-resolution data but also subcellular ST technologies (CosMx, MERFISH, and Xenium).

## Results

### Overview of SpaceBender Architecture

SpaceBender is a computational framework adapted off of the deep generative model CellBender^13^ which was originally designed to denoise single-cell data. The nature of ST data hinders the direct application of single-cell denoising methods; for example, RNA diffusion becomes spatially-dependent not uniform. In SpaceBender, we explicitly account for spatial-specific features by 1) leveraging automated tissue detection native to ST data to define empty observations for spot-resolution data and 2) defining ambient RNA profiles through local spatial niches (**Fig. 1A**, see **Methods**).

**Fig. 1:**
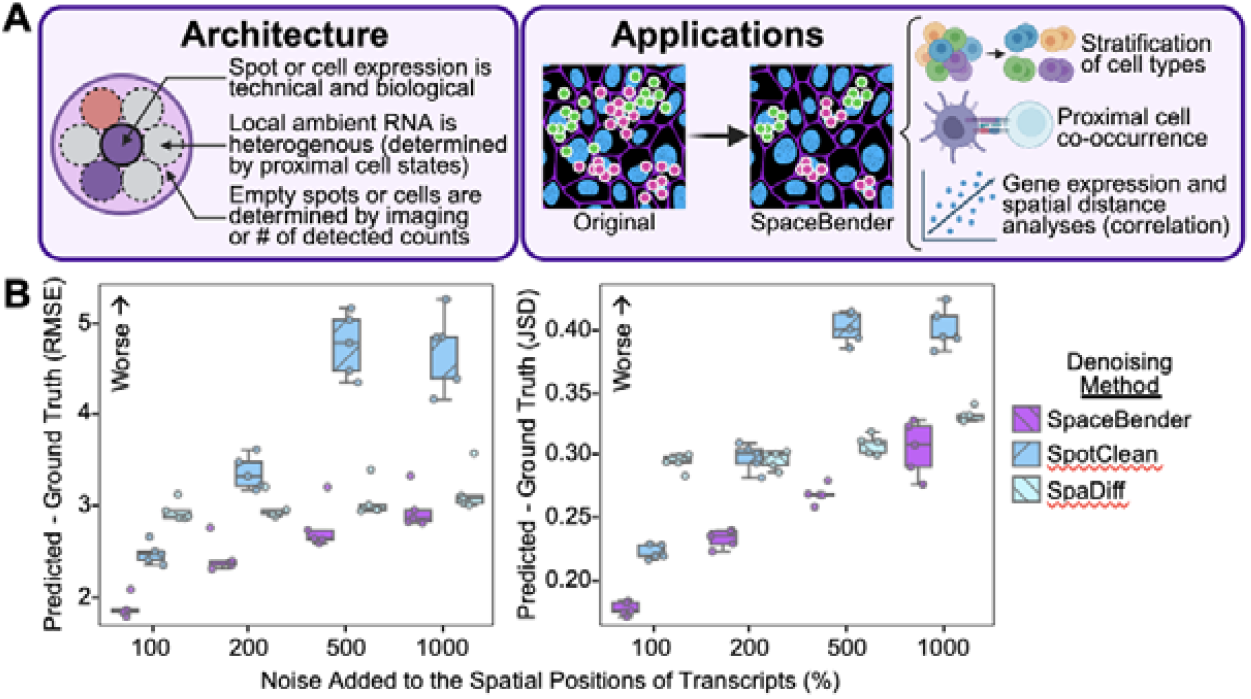
Architecture of and Benchmarking SpaceBender with Simulation Data. a) Depiction of the SpaceBender computational framework and example applications. b) Box plots of the denoising performance of SpaceBender (purple), SpotClean (blue), and SpaDiff (light blue) on simulated data with the x-axis representing the degree of noise added to the data (e.g. 100% = transcript positions were added to a uniform distribution of spot radius times 100%) and the y-axis representing the difference between the denoised matrix and the original data (quantified by root mean squared error [RMSE] and Jensen-Shannon divergence [JSD]). Box plots represent interquartile range (25^th^ to 75^th^ percentiles, IQR) with whiskers representing as 1.5 times IQR. Dots represent different simulated tissues.

### Benchmarking ST Denoising with Simulations

SpaceBender was benchmarked against current ST denoising methods (SpotClean^7^ and SpaDiff^10^) using simulated and, publicly-available, chimeric (mouse-human) ST data. Simulated data was generated in a manner consistent with previous works^10^ (see **Methods**). In brief, we simulated spot-resolution ST data with varying degrees of noise (RNA diffusion) by computationally aggregating subcellular-resolution ST data (Xenium, see **Data Availability**) using a grid of spots. Noise was defined as the addition of a uniform distribution to the spatial positions of transcripts; bounds of the distribution were set to the spot radius multiplied by the degree of noise (e.g. 100% noise = transcript positions can be moved up to a distance of radius times 100%). Denoising performance was quantified as the difference between the original and denoised spot-by-expression matrix; the metrics root-mean-squared-error (RMSE) and Jenson-Shannon-divergence (JSD) were utilized for these comparisons.

In these simulations, SpaceBender consistently outperformed current ST denoising methods at all noise levels (perturbation of transcript positions by 100-1000% spot radius) (**Fig. 1B**). For example, at a noise level of 100%, we observed an RMSE of 1.88±0.10 with SpaceBender, 2.47±0.10 with SpotClean, and 2.92±0.10 with SpaDiff (reported as mean ± 95% confidence interval, lower values indicate better performance). As expected, all methods performed worse with increasing degrees of noise (e.g. 1000% vs. 100%). We then asked if the denoising performance of SpaceBender was dependent on a specific set of model parameters. To this end, we varied the following key parameters of SpaceBender and interrogated for performance differences: radius size utilized to calculate spatial ambient RNA profiles, number of epochs for which SpaceBender was run, the model learning rate, number of samples taken from the posterior, and the number of dimensions utilized for the model embedding. These parameter variations, in vast majority, did not affect the denoising performance of SpaceBender (**Fig. S1**); as expected, increased learning rate, with the same number of epochs, decreased model performance due to the instability this parameter change introduces to model training.

### Benchmarking ST Denoising with Chimeric Tissues

To evaluate the performance of SpaceBender and current ST denoising methods on *in vivo* tissues, we leveraged publicly-available chimeric (mouse-human) ST data (Visium, 55µm spot diameter resolution)^7^. RNA diffusion can be formally defined in chimeric tissue as the degree of transcript bleed-over from one species’ tissue fragment into the other. These species-specific tissue fragments can be defined pathologically and computationally; both annotations were utilized for benchmarking to ensure model performance did not vary with minor annotation differences (**Fig. 2A, S2A**).

**Fig. 2:**
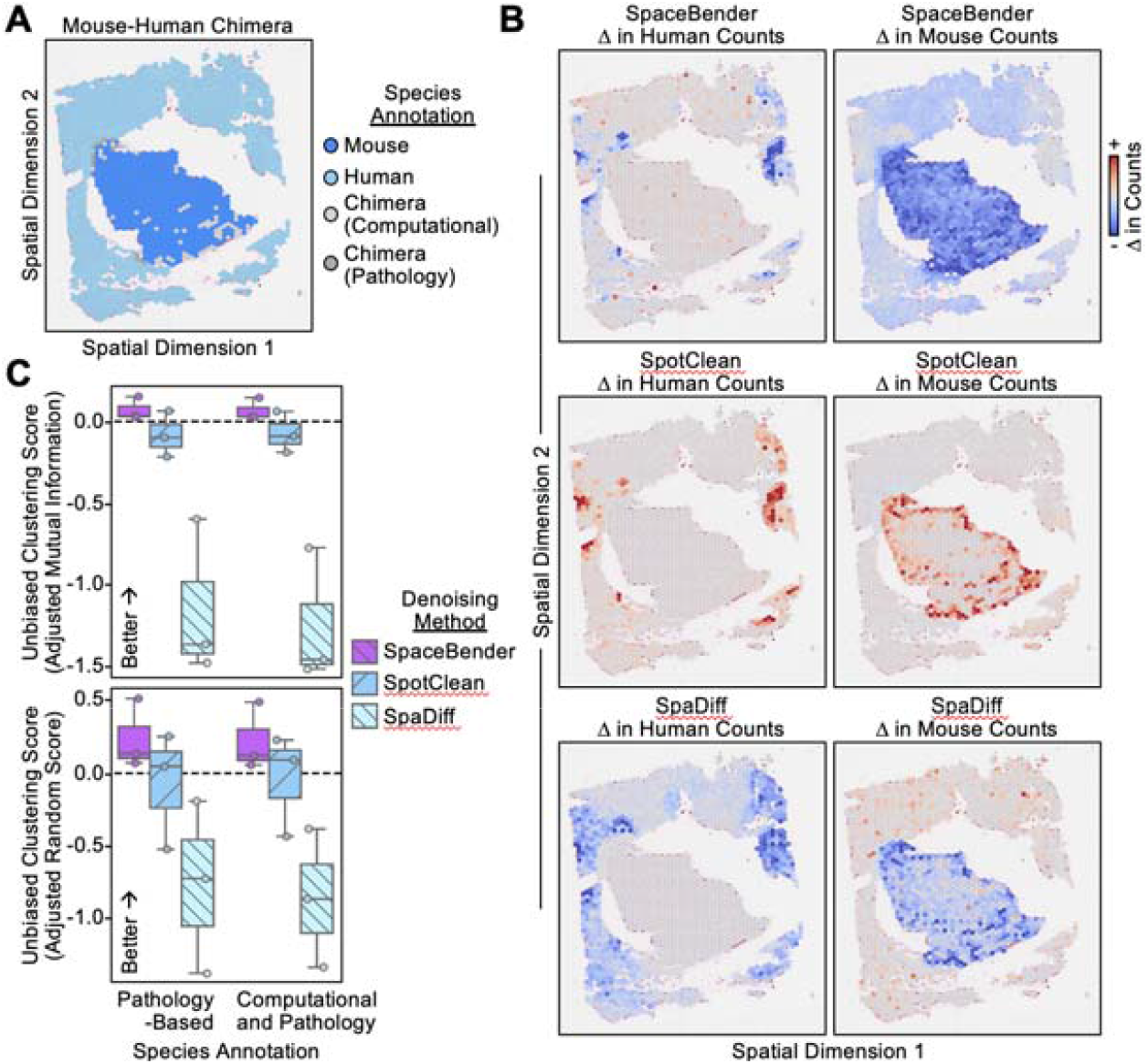
Benchmarking SpaceBender with Mouse-Human Chimeric Data. A) Spatial scatter plot of a representative chimeric (mouse-human) tissue with pathology and computationally derived annotations colored onto each dot; annotations were provided by the original authors of the publicly-available dataset. See legend on the right for color mapping of annotations. B) Spatial scatter plot of the changes in count number for genes from human (left) and mouse (right) genomes after processing by SpaceBender (upper), SpotClean (middle), or SpaDiff (lower). See color bars on the right for the mapping of the change in counts to spot color. C) Box plots of the denoising performance of SpaceBender (purple), SpotClean (blue), and SpaDiff (light blue) on chimeric tissues with the x-axis representing the annotations utilized (pathology or pathology and computationally derived) and the y-axis representing clustering evaluation metric scores (quantified by adjusted random index and adjusted mutual information). Box plots represent interquartile range (25th to 75th percentiles, IQR) with whiskers representing as 1.5 times IQR. Dots represent different tissues.

This analysis provided a visual example for denoising by SpaceBender; in this tissue, counts deemed as noise were removed from both intra- and inter-species fragments with an emphasis on tissue boundaries (between species) and an avoidance of a low cellularity region (upper border of the slide) (**Fig. 2B** upper). In an analogous but opposite manner, SpotClean also focused on tissue borders and avoided low cellularity regions but added, not removed, counts (**Fig. 2B** middle). SpaDiff, both added and removed counts, compared to the raw spot-by-expression matrix, and, although SpaDiff did focus on tissue borders, it appeared to add counts for mouse genes in the human tissue fragment (**Fig. 2B** lower).

ST denoising performance was quantified by both adjusted mutual information (AMI) and adjusted random score (ARS), in line with prior works^7,10^, and found SpaceBender to outperform current ST denoising methods (**Fig. 2C**); additional, less commonly used, metrics were also calculated and confirmed increased performance by SpaceBender compared to current methods (**Fig. S2B**). For example, we observed an AMI of 0.11±0.09 with SpaceBender, -0.04±0.09 with SpotClean, and -1.27±0.63 with SpaDiff using pathology and computational based species annotation (reported as mean ± 95% confidence interval, higher values indicate better performance). In all, through both simulations and *in vivo* chimeric tissue, we find SpaceBender to outperform existing methods in the denoising of ST data.

### ST Denoising Identifies Hidden Heterogeneity in Human Lymph Node

Although the performance of SpaceBender in simulations and chimeric tissue is promising, we sought to investigate if this method could increase the biological insights one could derive from ST data. To this end, we applied SpaceBender to a spot-resolution (Visium, 55µm spot diameter resolution) dataset of a human lymph node (see **Data Availability**); this tissue was chosen due to its marked spatial architecture (e.g. follicles, light and dark zones)^15^. Spot-resolution transcriptomes (original and SpaceBender processed) were subject to unbiased (Leiden) clustering and projected onto two dimensions, via uniform manifold approximation projection (UMAP). As expected, lymph node follicles could clearly be distinguished (Leiden 7 of the original ST data) (**Fig. 3A**). Intriguingly, Leiden clusters computed from SpaceBender processed data split the original Leiden 7 (follicles) into two subpopulations, SpaceBender Leidens 4 and 5 (**Fig. 3B**).

**Fig. 3:**
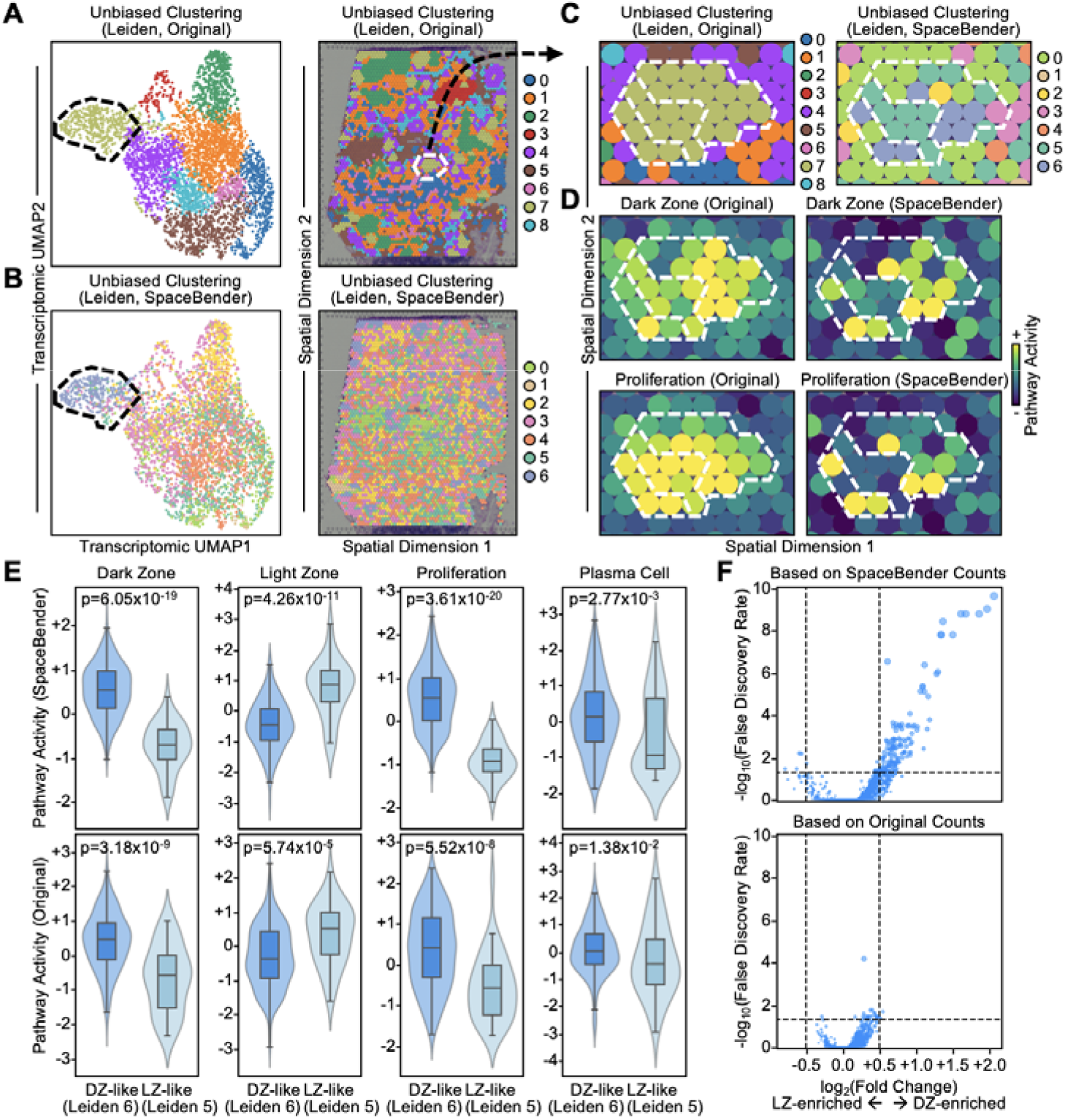
Denoising with SpaceBender Reveals Lymph Node Follicle Heterogeneity. A) Left: UMAP of spot transcriptomes from the original data with spot color representing unbiased (Leiden) cluster assignment based on the original transcriptomes. Right: Spatial scatter plot of spots colored by their Leiden cluster. See legend on the right for mapping between color and Leiden cluster. B) Left: UMAP of spot transcriptomes after SpaceBender processing with spot color representing unbiased (Leiden) cluster assignment based on the SpaceBender processed data. Right: Spatial scatter plot of spots colored by their Leiden cluster. See legend on the right for mapping between color and Leiden cluster. C) Spatial scatter plot of a follicle of interest with spots colored by their original (left) and SpaceBender (right) Leiden cluster assignments. See legends on the right for mapping between color and said Leiden clusters. White bounding box highlights LZ- and DZ-like regions corresponding to SpaceBender Leiden assignments. D) Spatial scatter plot of a follicle of interest with spots colored by pathway enrichment computed from the original (left) and SpaceBender processed (right) data. See color bar on the right for mapping between degree of enrichment and color. White bounding box highlights LZ- and DZ-like regions. E) Violin plots of the enrichment (y-axis) of various pathways in LZ-versus DZ-like regions (x-axis). Inner box plots represent interquartile range (25th to 75th percentiles, IQR) with whiskers representing as 1.5 times IQR. P-values were calculated using the Mann-Whitney U test. F) Scatter plot of the log_2_ fold change of gene expression in DZ-versus LZ-like regions (x-axis) against the statistical significance of said differences as quantified by -log_10_ False Discovery Rate (FDR, y-axis). P-values were calculated using the Mann-Whitney U test and subject to FDR multiple hypothesis testing correction.

We next sought to investigate if the original follicle cluster subdivisions, generated by SpaceBender processing, represent true biological heterogeneity. It is well known that lymph node follicles comprise of two distinct zones: B cell rich dark zones (DZ) that facilitate plasma cell differentiation post class switch recombination and light zones (LZ) where B cells interact with co-stimulatory T follicular helper and dendritic cells^15^. Thus, it may be that ST denoising allows for the identification of these zones within follicles. To interrogate for this, we first isolated a relatively larger follicle ascribed to only original Leiden 7 but SpaceBender Leidens 4 and 5 (**Fig. 3C, S3**). We next quantified the activity of canonical follicular-related pathways (e.g. DZ, proliferation) via pathway enrichment analysis (see **Methods**). This revealed SpaceBender Leidens 4 and 5 to correspond to LZ- and DZ-like regions, respectively (**Fig. 3D**). DZ pathway activity was more specific to the DZ-like region when calculated from the data denoised by SpaceBender. Further, only after SpaceBender processing could the, expected enrichment of proliferation pathways be observed in the DZ-like region.

To confirm that our findings were not specific to this singular follicle, we performed the aforementioned analysis on all follicles (original Leiden 7). Consistent with our prior analysis, LZ- and DZ-like regions were only separated after SpaceBender processing (**Fig. S4**). Notably, SpaceBender Leidens were not separated even when overlaid on the transcriptomic UMAP computed from the original ST data. This suggests that SpaceBender can separate of LZ-from DZ-like regions as a function of the model’s denoising rather than a technical artifact, e.g. a function of Leiden resolution (tunable parameter of clustering). Enrichment of LZ- and DZ-specific pathways was markedly more apparent in the SpaceBender processed data (**Fig. 3E**). For example, statistical testing (Mann-Whitney U test) of the difference in proliferation pathway enrichment between LZ- and DZ-like regions resulted in a p-value of 4.29×10^−5^ when computed from the original data but 4.11×10^−16^ when computed on SpaceBender processed data.

An increase in statistical significance after SpaceBender processing was also observed in whole transcriptome differential gene expression analysis of LZ-versus DZ-like regions: n=75 genes < FDR 0.05 after SpaceBender processing versus n=2 genes with the original data (**Fig. 3F, S5A**). We then sought to query if the genes differentially expressed (FDR < 0.05) after SpaceBender processing held biological meaning, via pathway enrichment analysis. Indeed, genes statistically over-expressed in the DZ-like region were enriched for DNA repair (non-homologous end joining) and leukocyte differentiation functions, both of which are well known to occur in the DZ to facilitate B cell class switch recombination and subsequent differentiation into plasma cells (**Fig. S5B**). In all, we demonstrate how SpaceBender processing can unveil hidden heterogeneity and increase the statistical significance of biologically relevant insights in human lymph node (Visium) ST data.

### ST Denoising Separates Clonal Subpopulations in Human Melanoma

In our previous work, we reported on a patient with cutaneous melanoma whose tumor presented with a spatially distinct subclone with chromosomal loss of a segment of chromosome 15 comprising the B2M gene locus^16^; this loss results in deficient MHC-I antigen presentation required for tumor cell recognition and clearance by cytotoxic T cells^17^. Although the majority of spots attributed to this subclone could be distinguished via decreased *B2M* expression, a clear delineation (spatially and transcriptomically) from other tumor spots was difficult due to the intermediate expression of *B2M* in spots along the subclone border, likely due to RNA diffusion and the ubiquitously high expression of B2M in nontumor cells. We investigated if SpaceBender, through denoising, is able to increase (“rescue”) the delineation of the B2M-loss subclone from other tumor cells.

To this end, said ST data (Visium) was processed via SpaceBender with malignant (tumor) spots isolated based on prior annotations^16^ (**Fig. S6A**). Consistent with our lymph node case study, SpaceBender processed data presented with greater statistical stratification of malignant versus non-malignant spots (evaluated via expression of melanoma marker genes, e.g. *MITF*^18^) (**Fig. S6B**). We then subject malignant spot transcriptomes, pre and post-SpaceBender processing, to unbiased (Leiden) clustering and two-dimensional projection; spots with decreased *B2M* expression were markedly more co-localized after SpaceBender processing (**Fig. 4A, S7A**). This held true whether the B2M-loss subclone was defined based on Leiden or individual spot expression of *B2M* (**Fig. 4B**). Across a range of *B2M* expression cutoffs, SpaceBender processing consistently allowed better separation of the B2M-loss subclone from other tumor spots in transcriptomic space (**Fig. 4C**). Further, consistent with the lymph node analyses, SpaceBender processing enhanced the statistical significance of differentially expressed genes called between the B2M-loss subclone and *B2M* expressing tumor spots (**Fig. 4D, S7B-C**). In all, through case studies of human lymph node and melanoma, we demonstrate the ability of SpaceBender to discover and enhance the biological insights that one can gleam from ST data.

**Fig. 4:**
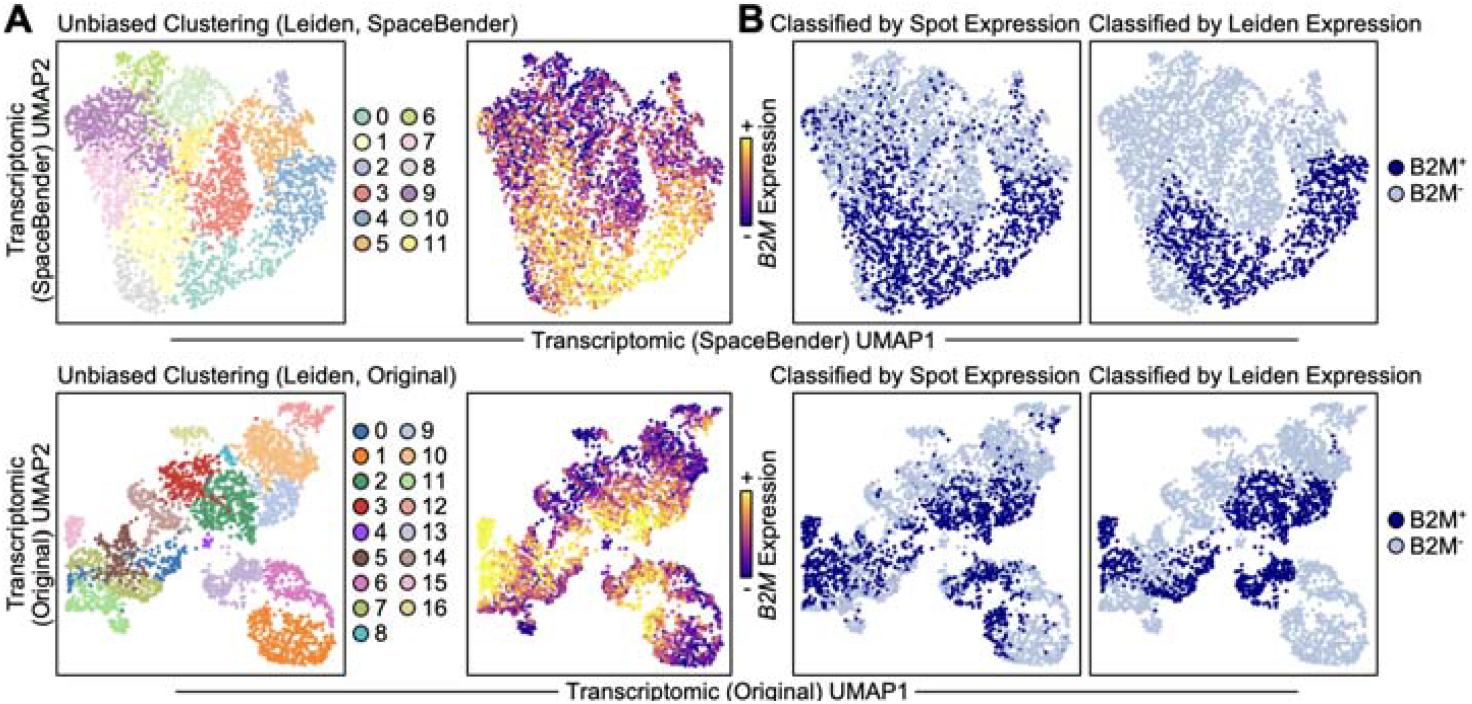
Denoising with SpaceBender Enhances the Separation of Melanoma Clone Transcriptomes. A) UMAP of spot transcriptomes from all malignant spots. UMAP was computed from the SpaceBender processed (upper) and original (lower) data. Spots are colored by their respective Leiden cluster assignment (left) or *B2M* expression (right). See legend on the right for mapping between color and Leiden cluster. See color bar on the right for mapping between color and *B2M* expression. B) UMAP of spot transcriptomes from panel (A) colored by B2M loss subclone status as defined at the resolution of individual spots (left) and Leiden clusters (right). C) Scatter plot of the goodness of clustering (quantified by Silhouette score, y-axis) of the B2M loss subclone versus other tumor spots at various *B2M* expression cutoffs used to define said clusters (x-axis). B2M loss subclone assignment was performed at the resolution of Leiden clusters (left) and individual spots (right). D) Violin plots of the change in Mann-Whitney U statistic in differentially upregulated (left) and downregulated (right) genes when comparing the whole transcriptomes of B2M loss subclone and other tumor spots. Inner box plots represent interquartile range (25th to 75th percentiles, IQR) with whiskers representing as 1.5 times IQR.

### Extension of SpaceBender to Subcellular Resolution ST Data

Currently available ST denoising methods, as well as the benchmarking and case studies presented above, focus on the cleaning of spot-resolution data (e.g. Visium). With the recent advent of subcellular ST technologies, there is now an additional need for the denoising of these subcellular resolution data (e.g. MERFISH, CosMx, Xenium). Thus, we sought to examine if SpaceBender could be extended to the denoising of said subcellular ST data.

To this end, we processed a MERFISH dataset of human tonsil using SpaceBender and subject the original and processed dataset to unbiased (Leiden) clustering and two-dimensional projection of single-cell transcriptomes (see **Methods**). SpaceBender processing noticeably increased the specificity of marker gene expression (**Fig. 5A-B**). For example, in the original data, B cell marker *CD79B*^19^ presented with high expression in B cell clusters (i.e. those co-expressing *FCER2*^20^ and *TNFRSF13C*^21^ [encoding BAFF-R]) and low-level non-specific expression in non-B-cell clusters (e.g. those expressing fibroblast marker *COL1A1*^22^). After SpaceBender processing, the expression of *CD79B* was restricted to B cell clusters. Similar denoising was observed in other cell type specific clusters, such as those enriched in T (*CD3D*^*+*^)^23^, myeloid (*CD14*^*+*^)^24^, proliferative (*MKI67*^*+*^)^25^, and endothelial (*PECAM1*^*+*^ [encoding CD31])^26^ cells. Denoising could be observed quantitatively through a decreased number of *CD3D*^*+*^*CD79B*^*+*^ cells (T-B doublets) after SpaceBender processing (**Fig. 5C**); these double-positive cells represent noise as *CD3D* and *CD79B* are mutually exclusive markers that define distinct cellular populations (T and B cells).

**Fig. 5:**
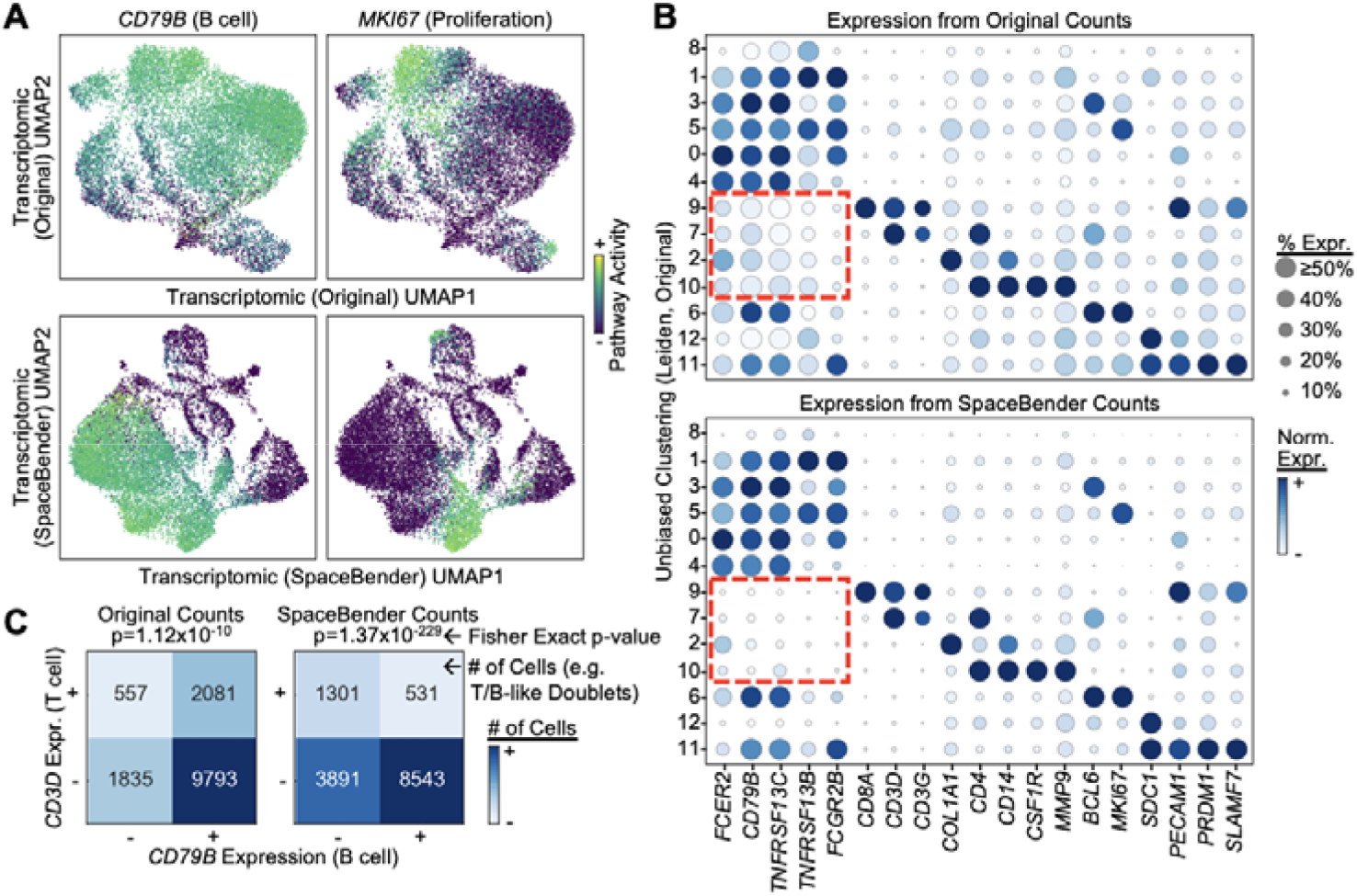
Extension of Denoising with SpaceBender to Subcellular Resolution Spatial Transcriptomics. A) UMAP of spot transcriptomes from MERFISH data of human tonsil. UMAP was computed from the original (upper) and SpaceBender processed (lower) data. Spots are colored by their respective expression of marker genes of interest. See legend on the right for mapping between color and Leiden cluster. B) Dot plots of the expression of marker genes of interest (columns) in each Leiden cluster (rows) as computed from the original (upper) and SpaceBender processed (lower) data. See legend on the right for mapping between dot size and percent of the given cluster that expresses said gene. See color bar on the right for mapping between color and expression of the given gene. Red bounding boxes denote genes of interest, corresponding to cluster (cell type) phenotypes, that change between original and SpaceBender processed data derived dot plots. C) Contingency tables of the number of cells expressing *CD3D* and/or *CD79B* in the original (left) and SpaceBender processed (right) data. Color represents the number of cells in a given quadrant, said number is annotated in each quadrant. P-values were computed from a Fisher’s exact test.

We repeated this denoising experiment with data from subcellular ST technologies CosMx (human non-small cell lung cancer [NSCLC]) and Xenium (human melanoma) (see **Data Availability**). Similar denoising results to the MERFISH data was observed in both datasets; non-specific marker gene expression was reduced after SpaceBender processing (**Fig. S8**). For example, MHC-II genes and *COL1A1* expression increased in specificity, became more cluster-specific, after SpaceBender processing in CosMx NSCLC dataset and immune cells no longer exhibited non-specific expression of melanocyte markers (e.g. *MITF*) in the Xenium melanoma dataset. In all, we demonstrate that SpaceBender can not only denoise spot-resolution ST data but can also be extended to the denoising of a myriad of subcellular ST technologies.

## Discussion

In this work, we presented SpaceBender as a method for ST denoising that outperforms currently available methods and, through several case studies, allows for the identification of new biological insights (e.g. LZ-versus DZ-like regions in **Fig. 3**) and statistical enhancement of these insights (e.g. increased log fold change and decreased FDR in **Figs. 3-4**). Denoised ST data synergizes with existing spatially informed methods, such as those for the detection of spatial-specific technical artifacts, identification of spatial niches, or quantification of spatially-restricted cell-cell communication^6,27–29^. By providing these tools with higher-quality clean data, their results become more accurate and thus, more informative, for example, due to fewer false positive or negative results.

The increasing availability of ST datasets, at increasingly larger scales, raises the enticing possibility of atlas-scale integration of ST data^30–33^. Single-cell atlases and integration efforts have already garnered numerous new biological insights, such as the identification of disease associated cell states and gene regulatory networks that may serve as potential therapeutic targets^34–37^. ST atlases would allow for the filtering of these insights for those that fit the spatial constraints of human tissue microenvironments, e.g. predicted interactions between cell types found to be spatially disparate likely represent false positive predictions or suggest that said cell types interact through secretory mechanisms. The development of ST atlases requires robust integration frameworks, which many works are pursuing^38^. These frameworks are likely to be enhanced by ST denoising methods, such as SpaceBender, which aid in the minimization of technical variation and thus, allow for true biological heterogeneity to be discovered. In all, SpaceBender serves to not only support the interrogation of specific biological questions through ST data, but we are hopeful that it may also aid current and future atlas-scale ST integration efforts that seek to distill generalizable and spatially-grounded therapeutic insights.

## Methods

### Model Architecture and Design

SpaceBender was designed upon the existing architecture of CellBender^13^ (see **Code Availability** for the new scripts and code changes contributed by this manuscript). The detailed additions are as follows. Ambient RNA profiles were adapted for spatial data by calculating (spatially) local rather than global profiles. Local profiles were calculated by taking the average expression of neighboring observations using k-nearest-neighbor (kNN) or distance-based identification of spatially-proximal neighbors. The decision to use the kNN- or distance-based method is made available to the user so they may choose the option that best fits their use case; distance-based methods were utilized in this manuscript. Additional changes involved the removal and adjustment of single-cell-specific parameters. For example, we have replaced the cell detection parameter with the default “in tissue” parameter provided by the Visium technology, a number-of-counts cutoff can also be utilized to determine which observations are “in tissue” although we have found SpaceBender to function without any cells marked as “out of tissue”. Detailed changes are made accessible in our **Code Availability** section where we provide access to the source code as well as the Python notebooks utilized for all presented analyses.

### Generation of and Benchmarking with Simulated Data

Simulated data were generated in a similar manner to prior works^10^. That is, publicly-available subcellular-resolution ST data (coronal slice of the mouse brain assayed with Xenium Prime, 10x Genomics) was computationally laid upon a spot-resolution (Visium-like) grid; transcripts were aggregated together if their spatial positions lay in the same computationally-defined spot. Noise was added to the simulated data by perturbing the spatial positions of each transcript, prior to spot-resolution aggregation, with a uniform distribution (minimum of zero, maximum of spot radius times noise level). For example, simulated data with a noise level of 100% means that the spatial positions of each transcript, prior to spot-resolution aggregation, was added to a uniform distribution with a minimum value of zero and maximum value equal to the radius of each spot.

This simulation process was performed n=5 randomly chosen fields-of-view (FOV); to evaluate ST denoising methods across multiple replicates. Simulated data from each FOV were generated with multiple degrees of noise (i.e. 100%, 200%, 500%, 1000%). Simulated data was then separately processed with SpotClean^7^, SpaDiff^10^, and SpaceBender (this manuscript). Denoising performance was quantified via root-mean-squared-error (RMSE) and Jensen-Shannon divergence (JSD) by comparing the expression matrices denoised by the aforementioned methods as compared to ground truth denoised expression matrices (prior to the computational addition of noise).

### Evaluating of Tunable Parameters on the Denoising Performance of SpaceBender

To investigate the impact of each tunable SpaceBender parameter on the method’s denoising performance, we repeated the aforementioned denoising study on simulated data with a range of values for each parameter (several folds above and below the default value). These parameters are the radius utilized to calculate the local spatial ambient RNA profile, number of epochs SpaceBender was trained for, the learning rate of this training process, the number of samples taken for posterior estimation, and the number of Z-dimensions (nodes in the hidden layer) utilized by the model. When a given parameter was being evaluated (tested at varying values), all other parameters were set to their default value to ensure we were only examining the effect of the tuned parameter.

### Benchmarking with Chimeric Data

Publicly-available mouse-human chimeric data^7^ was downloaded. Raw expression matrices were filtered to only include genes that were expressed in any (n≥1) cells. This chimeric data was processed in the same manner as the simulation data; that is, with SpotClean^7^, SpaDiff^10^, and SpaceBender (this work). Denoised expression matrices were normalized (log transform of library size normalized counts, ln(counts per 10K + 1)). These matrices were subject to unbiased (Leiden^39^, resolution=0.5) clustering after processing through principal component analysis (PCA) based dimension reduction and k-nearest-neighbors (kNN) graph calculation (using the aforementioned PCs as input). Species (mouse versus human) annotation were the same as the original authors’ annotations. Two sets of annotations are publicly-available, one that solely defines mouse-human border “mixture” tissue through pathology and another that also defined “mixture” tissue computationally. Denoising performance was evaluated using both sets of annotations to ensure that the results were not specific to one set of annotations. Clustering metrics (available from the Python package scikit-learn [sklearn]^40^) were utilized to evaluate denoising performance: adjusted random score, adjusted mutual information, homogeneity, completeness, V-measure, and Fowlkes-Mallows score. Unbiased (Leiden) clusters and mouse-human species annotations were compared in the aforementioned clustering metrics; roughly, these metrics quantify how well unbiased (Leiden) clusters separate spot-resolution observations by their mouse-human species annotations.

### Processing ST Data for Case Studies

Publicly available human lymph node (Visium), melanoma (Visium), and subcellular-resolution ST data (Xenium, CosMx, MERFISH) were downloaded and processed as follows, see Data Availability section for the exact links to said data. Raw counts were acquired from the aforementioned providers and filtered for genes that were expressed in any (n≥1) cells. Dataset expression matrices were then processed by SpaceBender, counts were normalized (log transformation of library size normalized counts, ln(counts per 10K + 1)), and then subject to dimension reduction. Dimension reduction was performed for both the original and SpaceBender processed expression matrices via PCA decomposition, kNN graph calculation of said PCs, unbiased (Leiden) clustering based on the kNN graph, (resolution of 0.5 was utilized for analysis of human lymph node and 1.0 for human melanoma and subcellular-resolution ST data), and two-dimensional projection of transcriptomes by UMAP^41^.

### Analysis of Human Lymph Node

Processed human lymph node data was scored for functional pathways by the “scanpy.tl.score_genes” function of the Scanpy package^42^; this function calculates pathway activity as the average expression of the genes of the pathway compared to the average expression of a control gene set (control genes are chosen as those with similar average expression, across all observations, as the genes in the given pathway). Pathways were retrieved from the Molecular Signatures Database (MSigDB)^43^; we focused on pathways relevant to the human lymph node (e.g. light-versus dark-zone regions of a lymph node follicle, lymphocyte proliferation). The plasma cell signature was calculated using the gene set of CD38^44^ and IRF4^45^, well known marker genes of plasma cells; CD138^44^ was not utilized due to its minimal expression across spots.

Observations (spots) that mapped to lymph node follicles (enrichment of follicle and germinal center related pathways) underwent an additional round of dimension reduction (PCA decomposition, kNN graph calculation on said PCs, UMAP calculation from the kNN graph). Light-(SpaceBender Leiden 5) and dark-zone (SpaceBender Leiden 6) like follicular regions (defined by their enrichments of light- and dark-zone related pathways from MSigDB, respectively) were compared statistically through the Mann-Whitney U test. Multiple hypothesis testing was corrected for by the Benjamin-Hochberg method (false discovery rate, FDR). Pathway enrichment analysis of differentially expressed genes (FDR < 0.05 and log_2_ of fold change) was performed via EnrichrR^46^.

### Analysis of Human Melanoma

Processed human melanoma data was subsetted for observations mapping to malignant cells; this annotation was derived from our prior work^16^. Subsetted data underwent an additional round of dimension reduction, similar to the human lymph node data (PCA decomposition, kNN graph calculation on said PCs, UMAP calculation from the kNN graph). *B2M* expression was normalized to Z-scores: the mean *B2M* expression was subtracted from the data and divided by the standard deviation of said expression (this was performed separately for the original and SpaceBender processed data). Observations (spot) were assigned as B2M^+^ or B2M^-^ by either normalized *B2M* expression of the given observation (spot-resolution assignment) or by the normalized *B2M* expression of the unbiased (Leiden) cluster the observation is a part of (Leiden- or cluster-resolution assignment).

The separation of spot- or (Leiden) cluster-assigned B2M^+^ and B2M^-^ observations in transcriptomic space (with or without SpaceBender processing), was evaluated via the Silhouette score. Spot-resolution assignments were performed at cutoffs from Z-scores of -0.25 to +0.75 at a step size of 0.1. Leiden- or cluster-resolution assignments were performed at cutoffs from Z-scores of -0.25 to +0.75 at a step size of 0.1; unbiased (Leiden) clustering was performed at multiple resolutions (i.e. 0.75, 1.00, 1.25, 1.50) to ensure results were not specific to one clustering of the data. B2M^+^ and B2M^-^ spots were statistically compared through the Mann-Whitney U tests. Multiple hypothesis testing was corrected for by the Benjamin-Hochberg method (FDR).

### Analysis of Subcellular-Resolution ST Data

Processed subcellular-resolution ST data from original and SpaceBender processed expression matrices were qualitatively compared by the expression of cell-type specific marker genes (e.g. *CD3D* for T cells^23^) across unbiased (Leiden) clusters. Original and SpaceBender processed expression matrices were quantitatively compared by “doublet” expression, that is cells that co-expressed marker genes known to solely be expressed in a singular cell-type (i.e. cells that co-expressed T cell specific marker *CD3D* and B cell specific marker *CD79B*^19^). The Fisher-Exact test was utilized to quantify the statistical significance of the contingency tables created by counting the number of cells with *CD3D* and/or *CD79B* expression (that is more than zero counts of said gene). This statistical test was chosen because it is the non-parametric (does not assume a statistical distribution) version of the Chi-squared test. Subcellular-resolution data were down-sampled to specific spatial regions, randomly chosen, due to the large size of said data; the specific areas that were sampled for the presented analyses are made available in the computational scripts provided in our Code Availability section.

### Quantification and Statistical Analysis

Statistical tests were performed using the Mann-Whitney U test, chosen due to its non-parameter nature (i.e. it does not assume the data to from a specific statistical distribution). P-values were corrected for multiple hypothesis testing via the Benjamin-Hochberg method (also known as false discovery rate, FDR).

## Data Availability

All data was publicly available. Xenium of mouse brain utilized for simulation studies was downloaded from https://www.10xgenomics.com/datasets/xenium-prime-fresh-frozen-mouse-brain. For case studies, Visium of human lymph node was downloaded from https://www.10xgenomics.com/datasets/human-lymph-node-1-standard-1-1-0, Visium of human melanoma was retrieved from our prior work^16^, MERFISH of human tonsils was downloaded from https://www.ncbi.nlm.nih.gov/geo/query/acc.cgi?acc=GSM8649279 (GSE282714), CosMx of human non-small cell lung cancer was downloaded from https://brukerspatialbiology.com/products/cosmx-spatial-molecular-imager/ffpe-dataset/nsclc-ffpe-dataset/, and Xenium of human melanoma was downloaded from https://www.10xgenomics.com/datasets/xenium-prime-ffpe-human-skin. The processed outputs have been deposited in Zenodo (reviewer-accessible link is as follows): https://zenodo.org/records/19458447?token=eyJhbGciOiJIUzUxMiJ9.eyJpZCI6Ijk5YmY5ZWYxLTc1OTctNDE3My04MmEyLTk4NzFjZGQxY2ZiNSIsImRhdGEiOnt9LCJyYW5kb20iOiJhMDBhMTBlNjVhNGQ3NWU0MzBiYmY0OGQxN2IyODhjNSJ9.q090CklPMl3OYlC1uQaSnx2bmZOD1iXtR50NBazCt9nX76v3T6qR5SN7xVhyBEYc8XlGvmK4Z4YjezTYX-E2Kw.

## Code Availability

SpaceBender is freely available at https://github.com/danielgchen/SpaceBender as a freely available public, Python, package; tutorial notebooks are available in the same repository. Code for reproduction of the analyses in this manuscript are available at https://github.com/danielgchen/reproducibility_spacebender.

## Acknowledgements

D.G.C. is supported by the UCLA-Caltech Medical Scientist Training Program. A.R. is supported by the Parker Institute for Cancer Immunotherapy (PICI), NIH grants R35 CA197633 and P01 CA168585, the Ressler Family Fund, and the support from Ken and Donna Schultz, Todd and Donna Jones, Robert (Bob) and Mary Jean Rumer, Karen and James Witemyre, and Thomas Stutz through the Jonsson Cancer Center Foundation, and Jonathan Isaacson through the Melanoma Research Foundation. K.M.C. is supported by the V Foundation Gil Nickel Melanoma Research Fellowship, the Parker Institute for Cancer Immunotherapy, and the Melanoma Research Alliance-Ressler Family Fund Young Investigator Award.

## Supplementary Figures

**Fig. S1:**
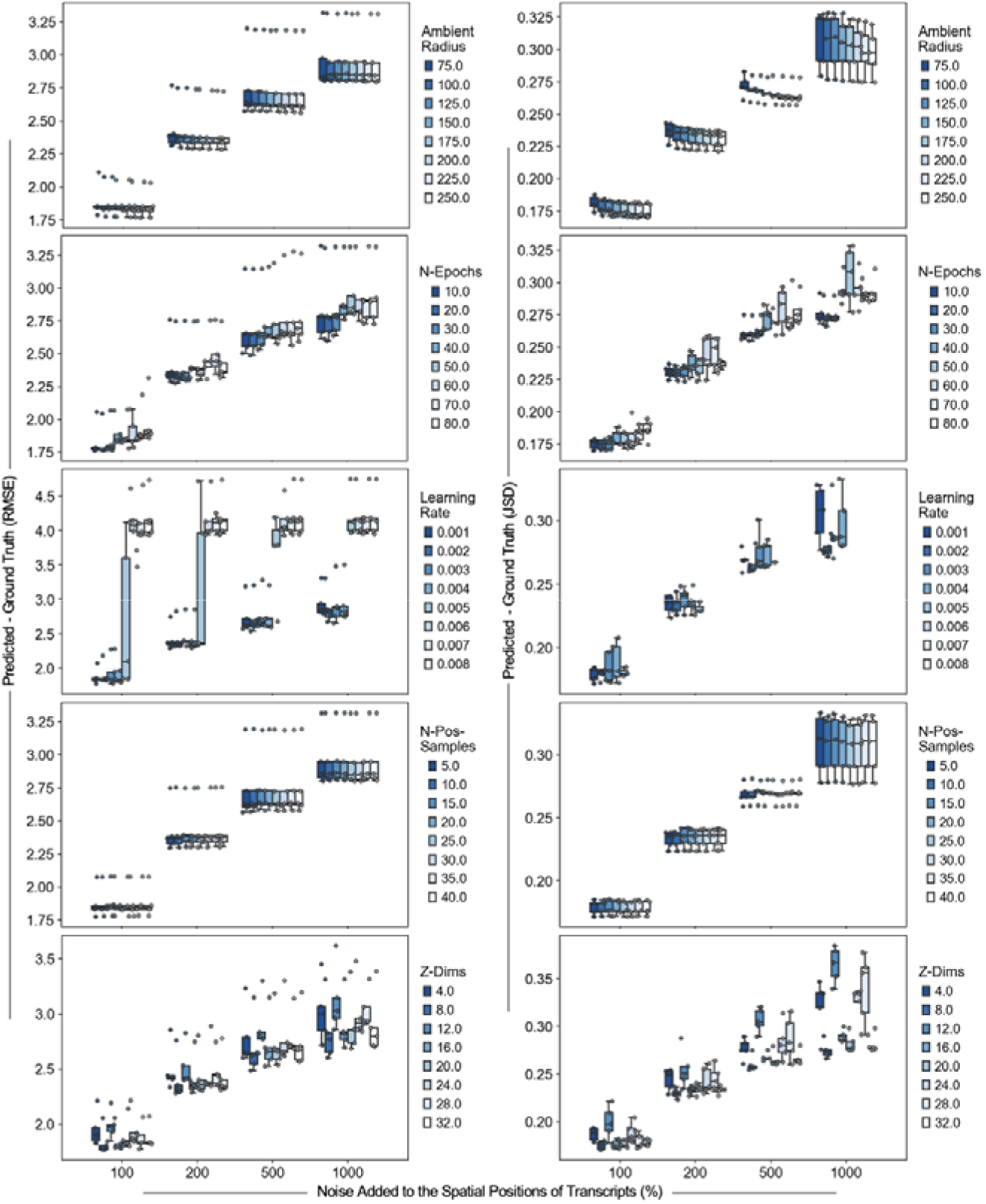
Impact of SpaceBender Parameters on Model Denoising Performance. Box plots of the denoising performance of SpaceBender on simulated data with the x-axis representing the degree of noise added to the data (e.g. 100% = transcript positions were added to a uniform distribution of spot radius times 100%) and the y-axis representing the difference between the denoised matrix and the original data (quantified by root mean squared error [RMSE] and Jensen-Shannon divergence [JSD]). Each panel represents a different model parameter being varied. See legend on the right for the mapping between box color and the varied parameter value. Box plots represent interquartile range (25th to 75th percentiles, IQR) with whiskers representing as 1.5 times IQR. Dots represent different simulated tissues.

**Fig. S2:**
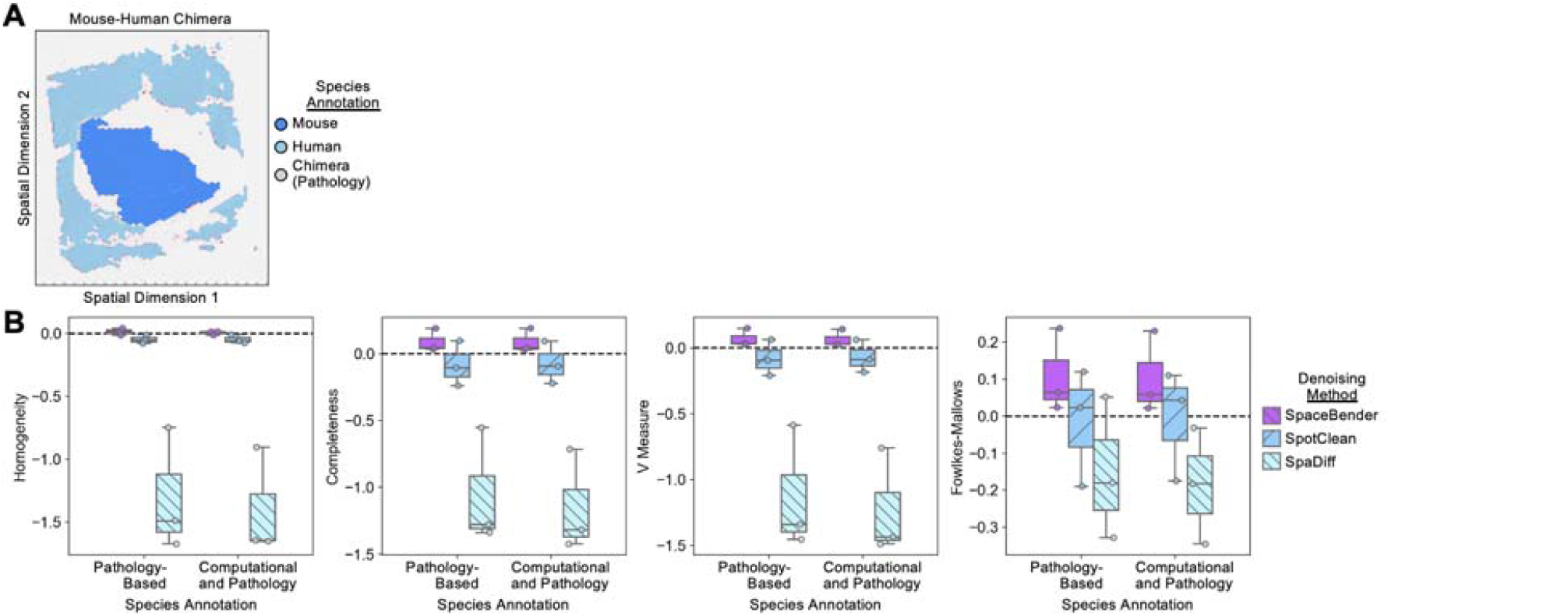
Extended Benchmarking Metrics of SpaceBender with Mouse-Human Chimeric Data. A) Spatial scatter plot of a representative chimeric (mouse-human) tissue with pathology derived annotations colored onto each dot; annotations were provided by the original authors of the publicly-available dataset. See legend on the right for color mapping of annotations. B) Box plots of the denoising performance of SpaceBender (purple), SpotClean (blue), and SpaDiff (light blue) on chimeric tissues with the x-axis representing the annotations utilized (pathology or pathology and computationally derived) and the y-axis representing clustering evaluation metric scores. Box plots represent interquartile range (25th to 75th percentiles, IQR) with whiskers representing as 1.5 times IQR. Dots represent different tissues.

**Fig. S3:**
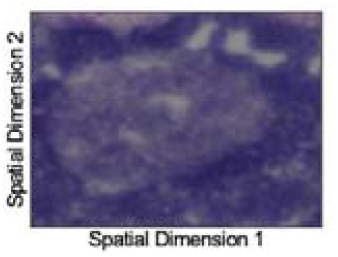
Imaging of a Lymph Node Follicle of Interest. Hematoxylin and eosin staining of the follicle of interest.

**Fig. S4:**
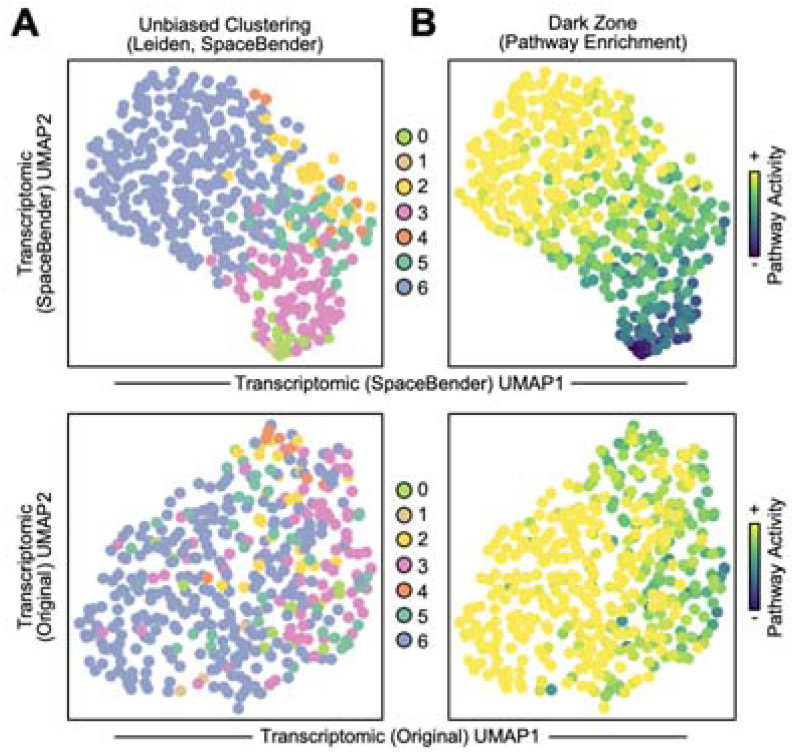
Dark- and Light-Zone Follicular Regions in the Lymph Node. A) UMAP of spot transcriptomes from all follicles in the given tissue (original Leiden 7). UMAP was computed from the SpaceBender processed (upper) and original (lower) data. Spots are colored by their SpaceBender Leiden cluster assignment. See legend on the right for mapping between color and Leiden cluster. B) UMAP of spot transcriptomes from panel (A) colored by their enrichment for DZ-related pathways. See color bar on the right for mapping between degree of enrichment and color.

**Fig. S5:**
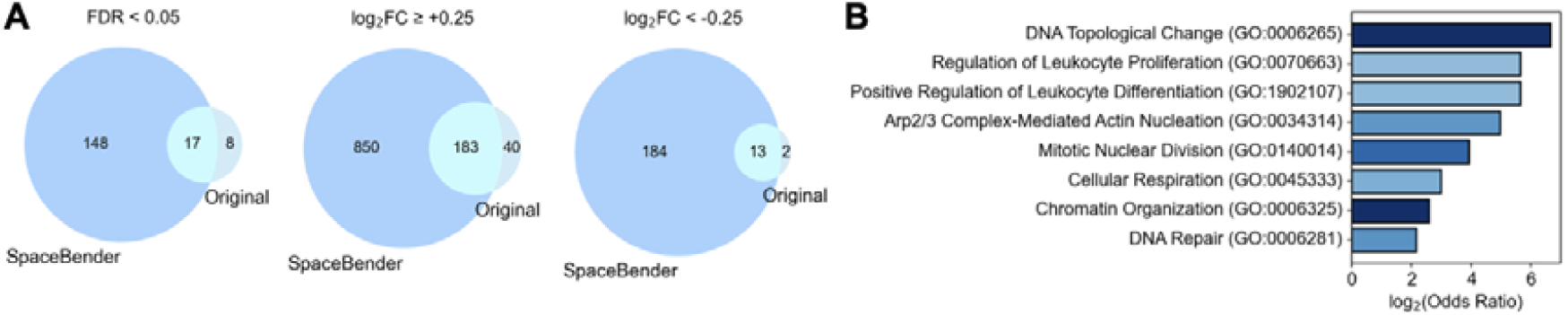
Denoising with SpaceBender Highlights Dark-Zone Specific Pathways. A) Venn diagrams between the differentially expressed genes identified in the original and SpaceBender processed data considering False Discovery Rate (FDR) and log_2_ fold change of gene expression between DZ- and LZ-like regions. B) Bar plot of the pathway enrichment of genes differentially upregulated in DZ-like regions after SpaceBender processing; all presented pathways are statistically significant (FDR < 0.05). Color represents statistical significance. See color bar on the right for mapping between degree of significance and color.

**Fig. S6:**
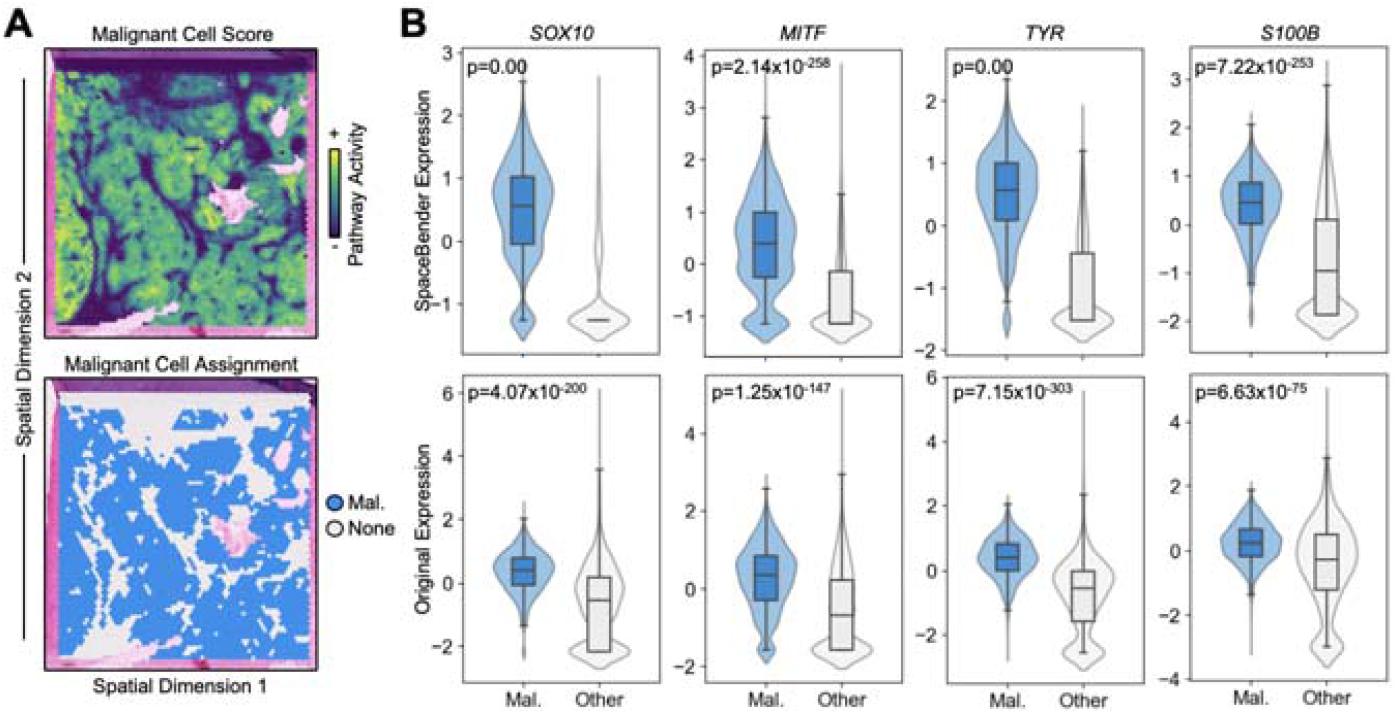
Denoising with SpaceBender Enhances the Separation of Malignant Melanoma Spot Transcriptomes from Adjacent Tissue. A) Spatial scatter plot of a tumor biopsy from a patient with melanoma. Spots are colored by their malignancy status via a continuous score (upper) and binary assignment (lower); annotations were derived from our prior work^16^. B) Violin plots of the enrichment (y-axis) of various pathways in malignant versus non-malignant spots (x-axis). Inner box plots represent interquartile range (25th to 75th percentiles, IQR) with whiskers representing as 1.5 times IQR. P-values were calculated using the Mann-Whitney U test.

**Fig. S7:**
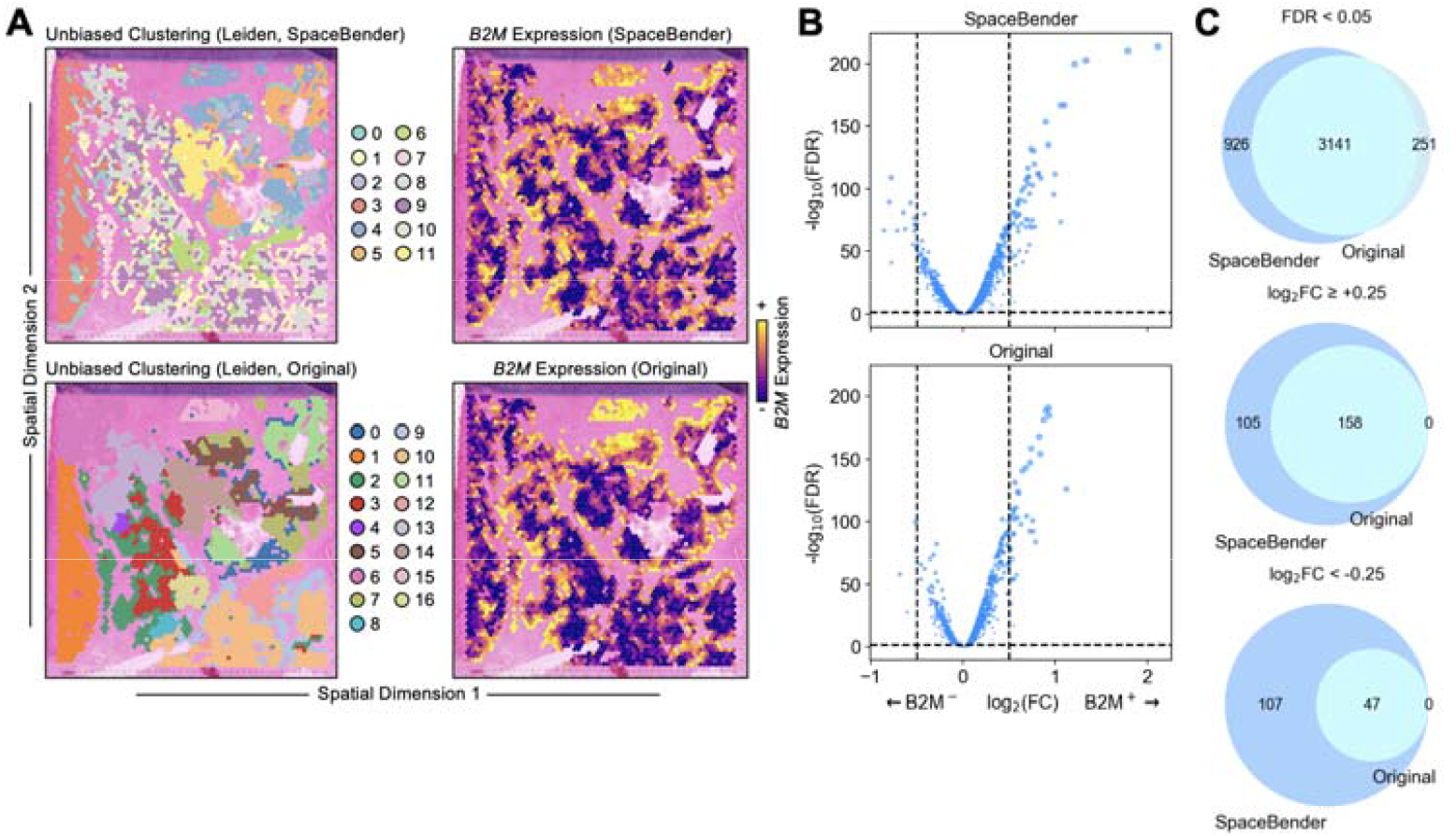
Denoising with SpaceBender Increases Statistical Differences in Gene Expression between Malignant Melanoma Clones. A) Left: Spatial scatter plot tumor spots from the tumor biopsy of a patient with melanoma. Spots are colored by their SpaceBender processed (upper) or original (lower) Leiden cluster assignment. Right: Spots are colored by their *B2M* expression. See legend on the right for mapping between color and Leiden cluster. See color bar on the right for mapping between *B2M* expression and color. B) Scatter plot of the log_2_ fold change of gene expression in B2M loss subclone versus other tumor spots (x-axis) against the statistical significance of said differences as quantified by -log_10_ False Discovery Rate (FDR, y-axis). P-values were calculated using the Mann-Whitney U test and subject to FDR multiple hypothesis testing correction. C) Venn diagrams between the differentially expressed genes identified in the original and SpaceBender processed data considering False Discovery Rate (FDR) and log_2_ fold change of gene expression between B2M loss subclone and other tumor spots.

**Fig. S8:**
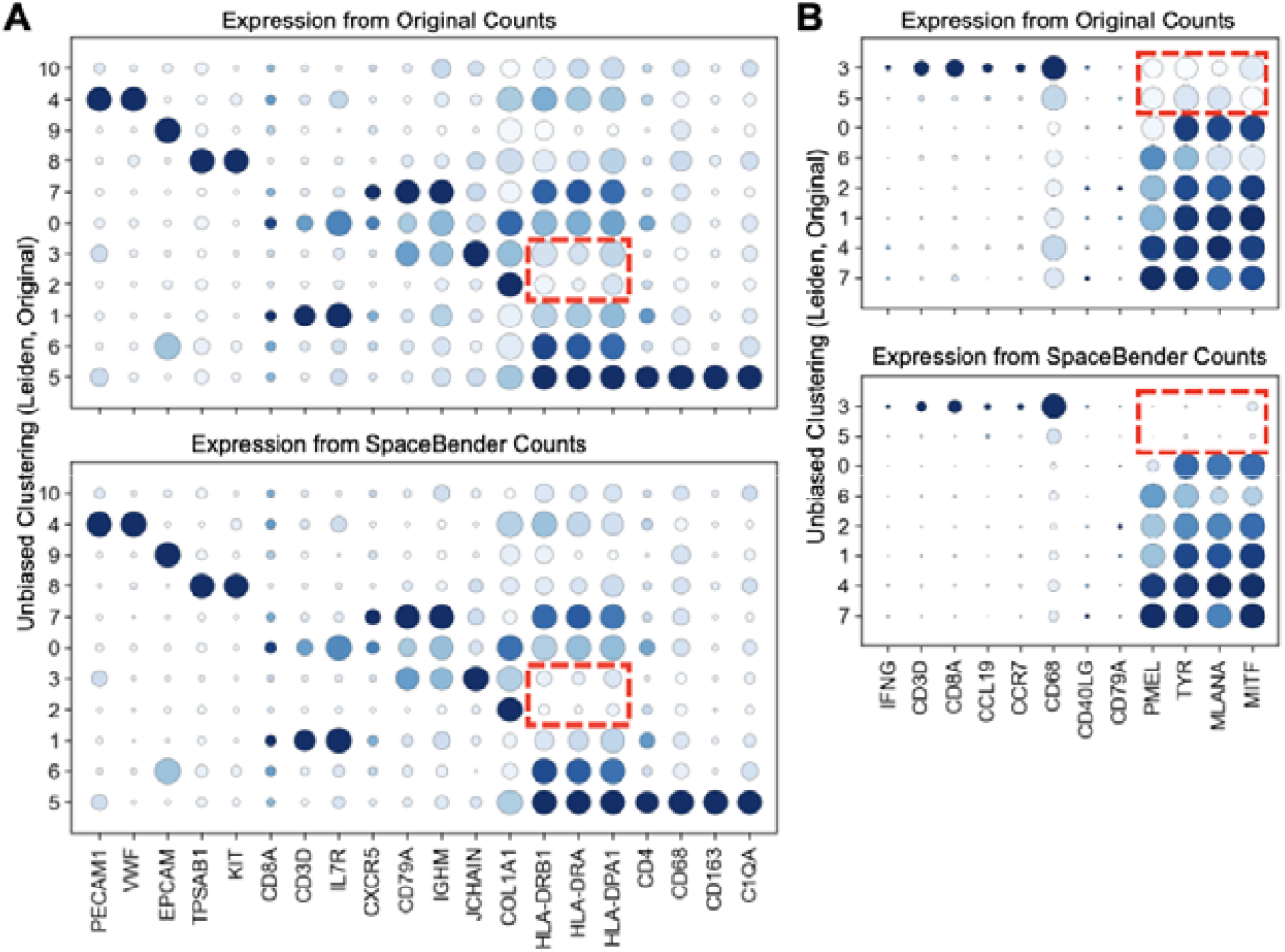
Extension of Denoising with SpaceBender to Xenium and CosMx. Dot plots of the expression of marker genes of interest (columns) in each Leiden cluster (rows) as computed from the original (upper) and SpaceBender processed (lower) CosMx non-small cell lung cancer (NSCLC) (panel [A]) and Xenium melanoma (panel [B]) data. See legend on the right for mapping between dot size and percent of the given cluster that expresses said gene. See color bar on the right for mapping between color and expression of the given gene. Red bounding boxes denote genes of interest, corresponding to cluster (cell type) phenotypes, that change between original and SpaceBender processed data derived dot plots.

