## Supplemental Figures for "SpaceBender: Denoising Spatial Transcriptomics Data to Enhance Biological Signals"

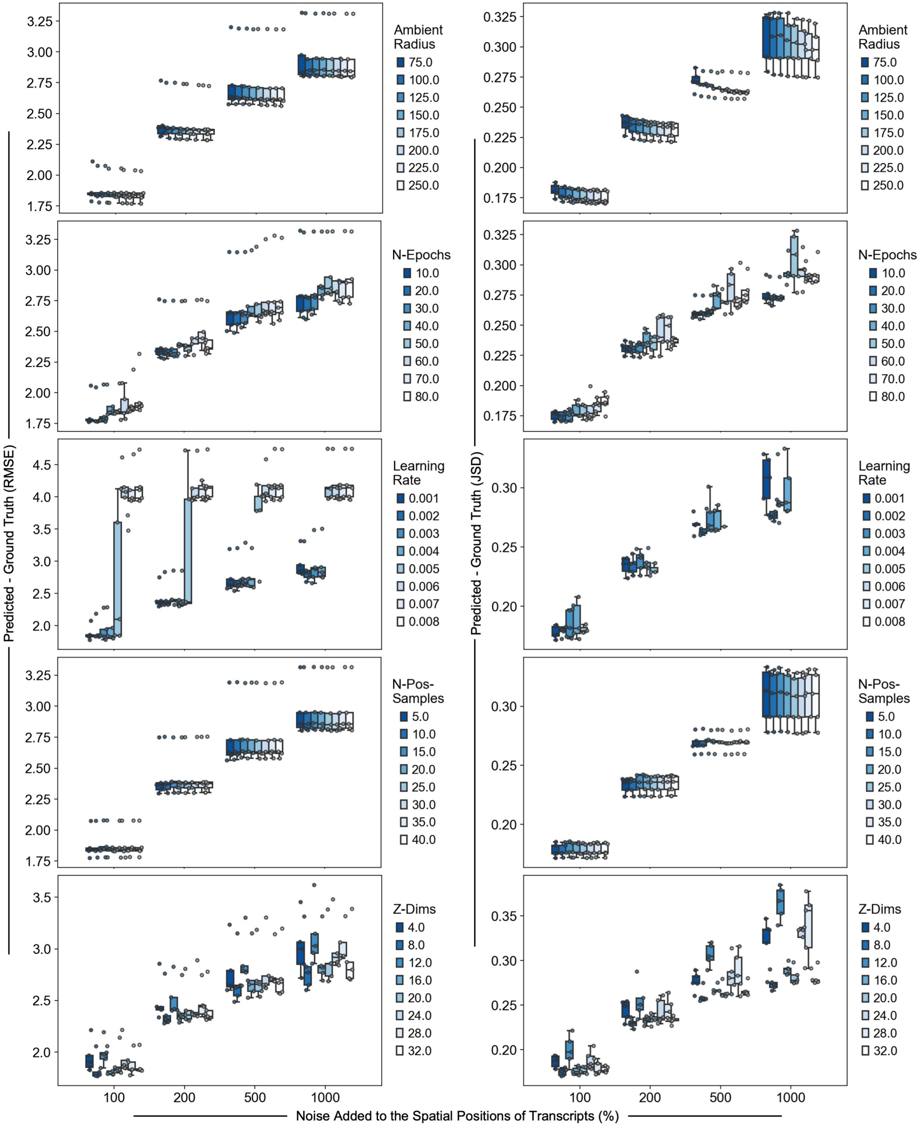


### Supplementary Figure 1

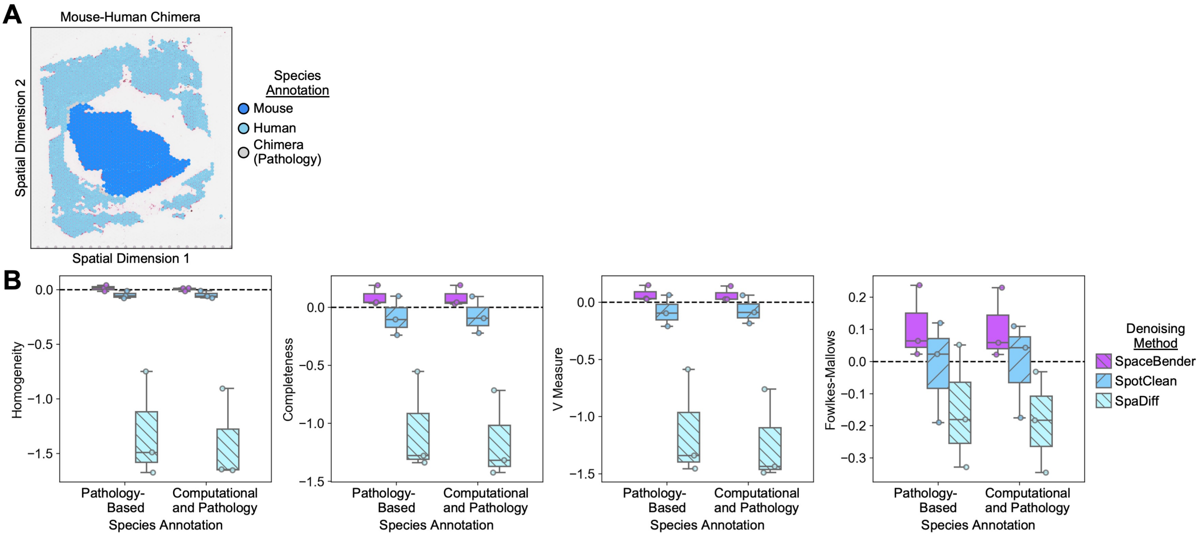


### Supplementary Figure 2

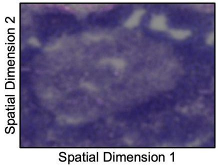


**Supplementary Figure 3**

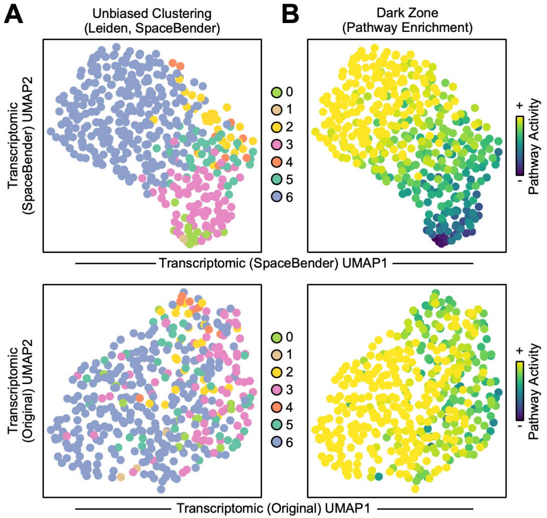


### Supplementary Figure 4

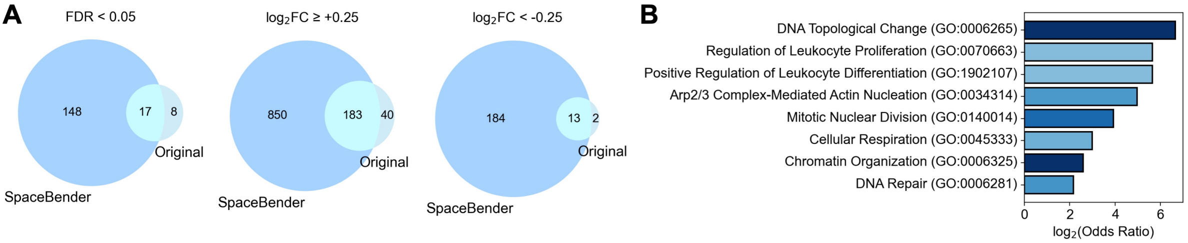


### Supplementary Figure 5

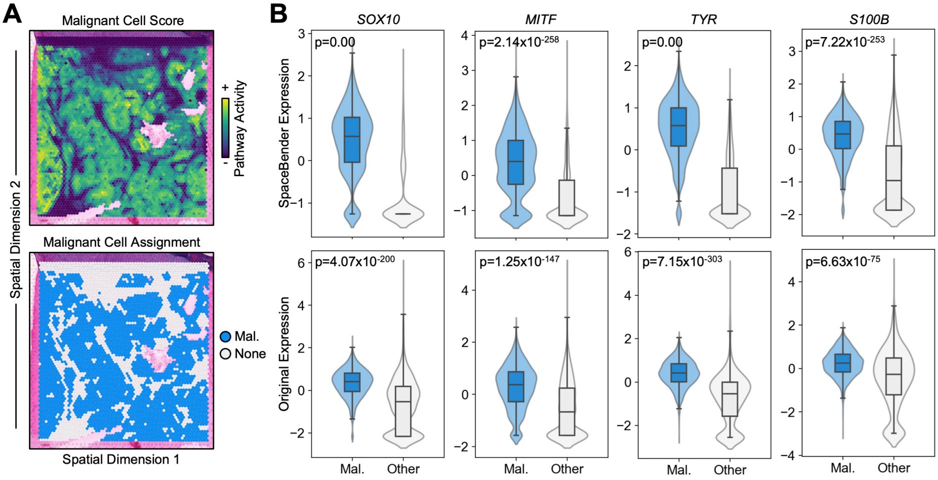


### Supplementary Figure 6

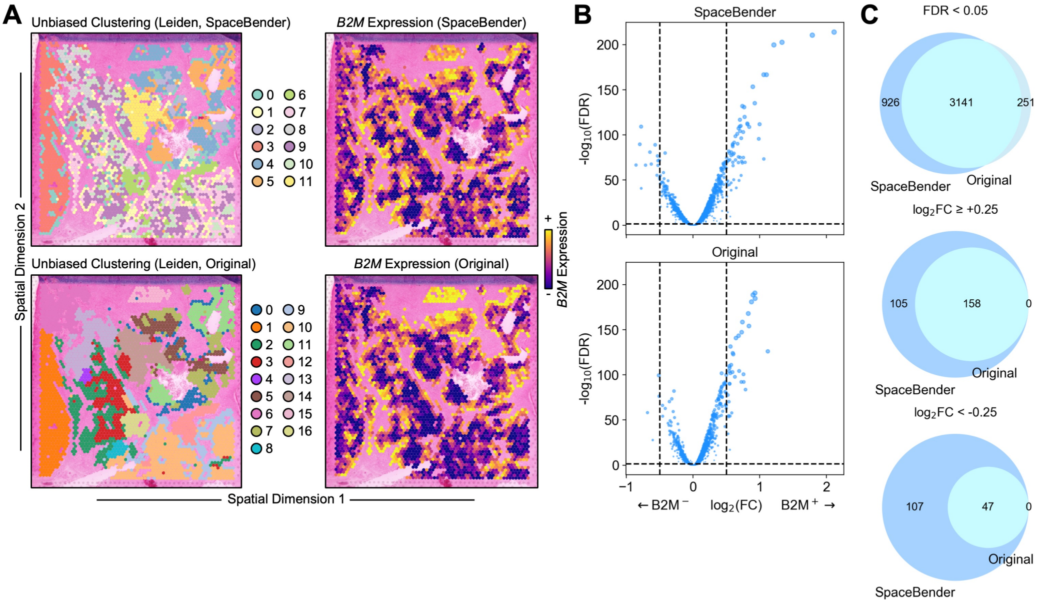


### Supplementary Figure 7

**A**

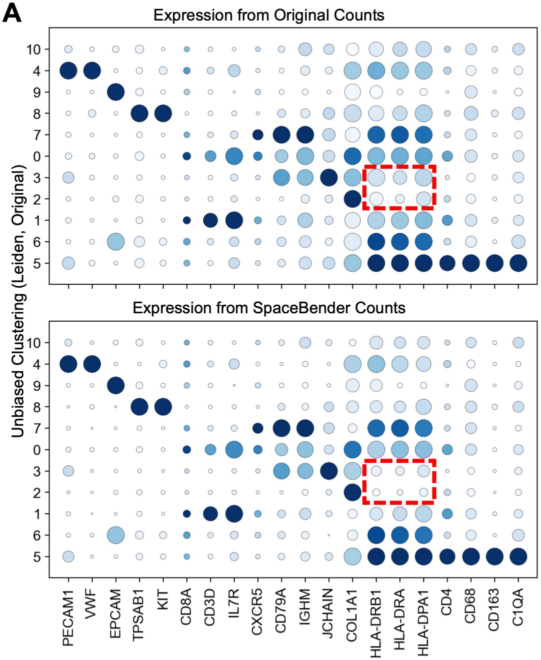


**B**

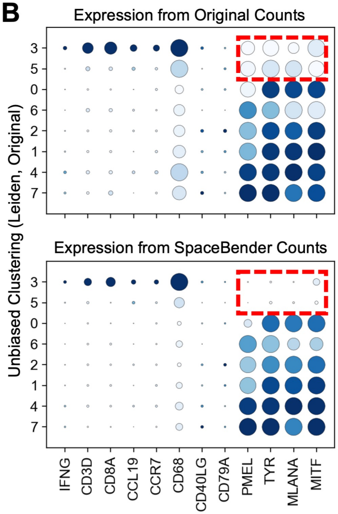


### Supplementary Figure 8
